# The influence of object-colour knowledge on emerging object representations in the brain

**DOI:** 10.1101/533513

**Authors:** Lina Teichmann, Genevieve L. Quek, Amanda K. Robinson, Tijl Grootswagers, Thomas A. Carlson, Anina N. Rich

**Affiliations:** Perception in Action Research Centre & Department of Cognitive Science, Macquarie University, Sydney, Australia; ARC Centre of Excellence in Cognition & its Disorders, Macquarie University, Sydney, Australia; School of Psychology, University of Sydney, Australia; Centre for Elite Performance, Training and Expertise, Macquarie University, Sydney, Australia; Donders Institute for Brain, Cognition and Behaviour, Radboud University, Nijmegen, The Netherlands

## Abstract

The ability to rapidly and accurately recognise complex objects is a crucial function of the human visual system. To recognise an object, we need to bind incoming visual features such as colour and form together into cohesive neural representations and integrate these with our pre-existing knowledge about the world. For some objects, typical colour is a central feature for recognition; for example, a banana is typically yellow. Here, we applied multivariate pattern analysis on time-resolved neuroimaging (magnetoencephalography) data to examine how object-colour knowledge affects emerging object representations over time. Our results from 20 participants (11 female) show that the typicality of object-colour combinations influences object representations, although not at the initial stages of object and colour processing. We find evidence that colour decoding peaks later for atypical object-colour combinations in comparison to typical object-colour combinations, illustrating the interplay between processing incoming object features and stored object-knowledge. Taken together, these results provide new insights into the integration of incoming visual information with existing conceptual object knowledge.

**Significance Statement:** To recognise objects, we have to be able to bind object features such as colour and shape into one coherent representation and compare it to stored object knowledge. The magnetoencephalography data presented here provide novel insights about the integration of incoming visual information with our knowledge about the world. Using colour as a model to understand the interaction between seeing and knowing, we show that there is a unique pattern of brain activity for congruently coloured objects (e.g., a yellow banana) relative to incongruently coloured objects (e.g., a red banana). This effect of object-colour knowledge only occurs after single object features are processed, demonstrating that conceptual knowledge is accessed relatively late in the visual processing hierarchy.

## Introduction

Successful object recognition depends critically on comparing incoming perceptual information with existing internal representations (Albright, 2012; Clarke & Tyler, 2015). A central feature of many objects is colour, which can be a highly informative cue in visual object processing (Rosenthal et al., 2018). Although we know a lot about colour perception itself, comparatively less is known about how object-colour knowledge interacts with colour perception and object processing. Here, we measure brain activity with magnetoencephalography (MEG) and apply multivariate pattern analyses (MVPA) to test how stored object-colour knowledge influences emerging object representations over time.

Colour plays a critical role in visual processing by facilitating scene and object recognition (Gegenfurtner & Rieger, 2000; Tanaka et al., 2001), and by giving an indication of whether an object is relevant for behaviour (Conway, 2018; Rosenthal et al., 2018). Objects that include colour as a strong defining feature have been shown to activate representations of associated colours (Bannert & Bartels, 2013; Hansen et al., 2006; Olkkonen et al., 2008; Teichmann et al., 2019; Vandenbroucke et al., 2014; Witzel et al., 2011), leading to slower recognition when there is conflicting colour information (e.g., a red banana; Nagai & Yokosawa, 2003; Tanaka & Presnell, 1999; for a meta-analysis, see Bramão, Reis, Petersson, & Faísca, 2011). Neuroimaging studies on humans and non-human primates have shown that there are several colour-selective regions along the visual ventral pathway (Lafer-Sousa et al., 2016; Lafer-Sousa & Conway, 2013; Seymour et al., 2010, 2015; Zeki & Marini, 1998). While the more posterior colour-selective regions do not show a shape bias, the anterior colour-selective regions do (Lafer-Sousa et al., 2016), supporting suggestions that colour knowledge is represented in regions associated with higher-level visual processing (Simmons et al., 2007; Tanaka et al., 2001). A candidate region for the integration of stored knowledge and incoming visual information is the anterior temporal lobe (ATL; Chiou et al., 2014; Papinutto et al., 2016; Patterson et al., 2007). In one study (Coutanche & Thompson-Schill, 2014), for example, brain activation patterns evoked by recalling a known object’s colour and its shape could be distinguished in a subset of brain areas that have been associated with perceiving those features, namely V4 and lateral occipital cortex, respectively. In contrast, recalling an object’s particular conjunction of colour *and* shape, could only be distinguished in the ATL, suggesting that the ATL processes conceptual object representations.

Time-resolved data measured with electroencephalography (EEG) or MEG can give an understanding of how conceptual-level processing interacts dynamically with perception. Previous EEG studies have examined the temporal dynamics of object-colour knowledge as an index of the integration of incoming visual information and prior knowledge (Lloyd-Jones et al., 2012; Lu et al., 2010; Proverbio et al., 2004). For example, Lloyd-Jones et al. (2012) showed that shape information modulates neural responses at ∼170ms (component N1), the combination of shape and colour affected the signal at 225ms (component P2), and the typicality of object-colour pairing modulated components approximately 225 and 350ms after stimulus onset (P2 and P3). These findings suggest that the initial stages of object recognition may be driven by shape, with the interactions with object-colour knowledge coming into play at a much later stage, perhaps as late as during response selection.

Using multivariate methods for time-resolved neuroimaging data, we can move beyond averaged measures (i.e., components) to infer what type of information is contained in the neural signal on a trial-to-trial basis. In the present study, we used MVPA to determine the timepoint at which neural activity evoked by congruently (e.g., yellow banana) and incongruently (e.g., red banana) coloured objects differs, which indicates when stored knowledge is integrated with incoming visual information. Furthermore, we examined whether existing knowledge about an object’s colour influences perceptual processing of surface colour and object identity. Overall, using colour as a model, our findings elucidate the timecourse of interactions between incoming visual information and prior knowledge in the brain.

## Materials and Methods

### Participants

20 healthy volunteers (11 female, mean age = 28.9 years, SD = 6.9 years, 1 left-handed) participated in the study. All participants reported accurate colour-vision and had normal or corrected-to-normal visual acuity. Participants gave informed consent before the experiment started and were financially compensated. The study was approved by the Macquarie University Human Research Ethics Committee.

### Stimuli

We identified five real world objects that previous studies have shown to be strongly associated with each of four different colours (red, green, orange and yellow; see Figure 1) (Bannert & Bartels, 2013; Joseph, 1997; Lloyd-Jones et al., 2012; Naor-Raz et al., 2003; Tanaka & Presnell, 1999; Therriault et al., 2009). Each colour category had one manmade object (e.g., fire hydrant), one living object (e.g., ladybird), and three fruits or vegetables (e.g., strawberry, tomato, cherry). We sourced two exemplar images for each object class, resulting in 10 images for each colour, 40 individual images in total. We then created incongruently coloured objects by swapping the colours (e.g., yellow strawberry, red banana). For both congruent and incongruent stimuli, we did not use the native colours from the images themselves, but instead overlayed pre-specified hues on desaturated (greyscale) images that were equated for luminance using the SHINE toolbox (Willenbockel et al., 2010). A greyscale image overlayed with its canonically associated colour (e.g., yellow hue applied to greyscale banana) resulted in a congruent object; a greyscale image overlayed with a colour different from its canonically associated colour (e.g., red hue applied to greyscale banana) resulted in an incongruent object. Every congruent object exemplar had a single colour-matched incongruent partner. For example, we used a specific shade of red and added it to the grey-scale images of the strawberry to make the congruent strawberry and overlayed it onto the lemon to make the incongruent lemon. We then took a specific shade of yellow and overlayed it on the lemons to make the congruent lemon exemplar, and onto the strawberry to make the incongruent strawberry exemplar. That means, overall, we have the identical objects and colours in the congruent and the incongruent condition, a factor that is crucial to ensure our results cannot be explained by features other than colour congruency. The only difference between these key conditions is that the colour-object combination is either typical (congruent) or atypical (incongruent).

**Figure 1.**
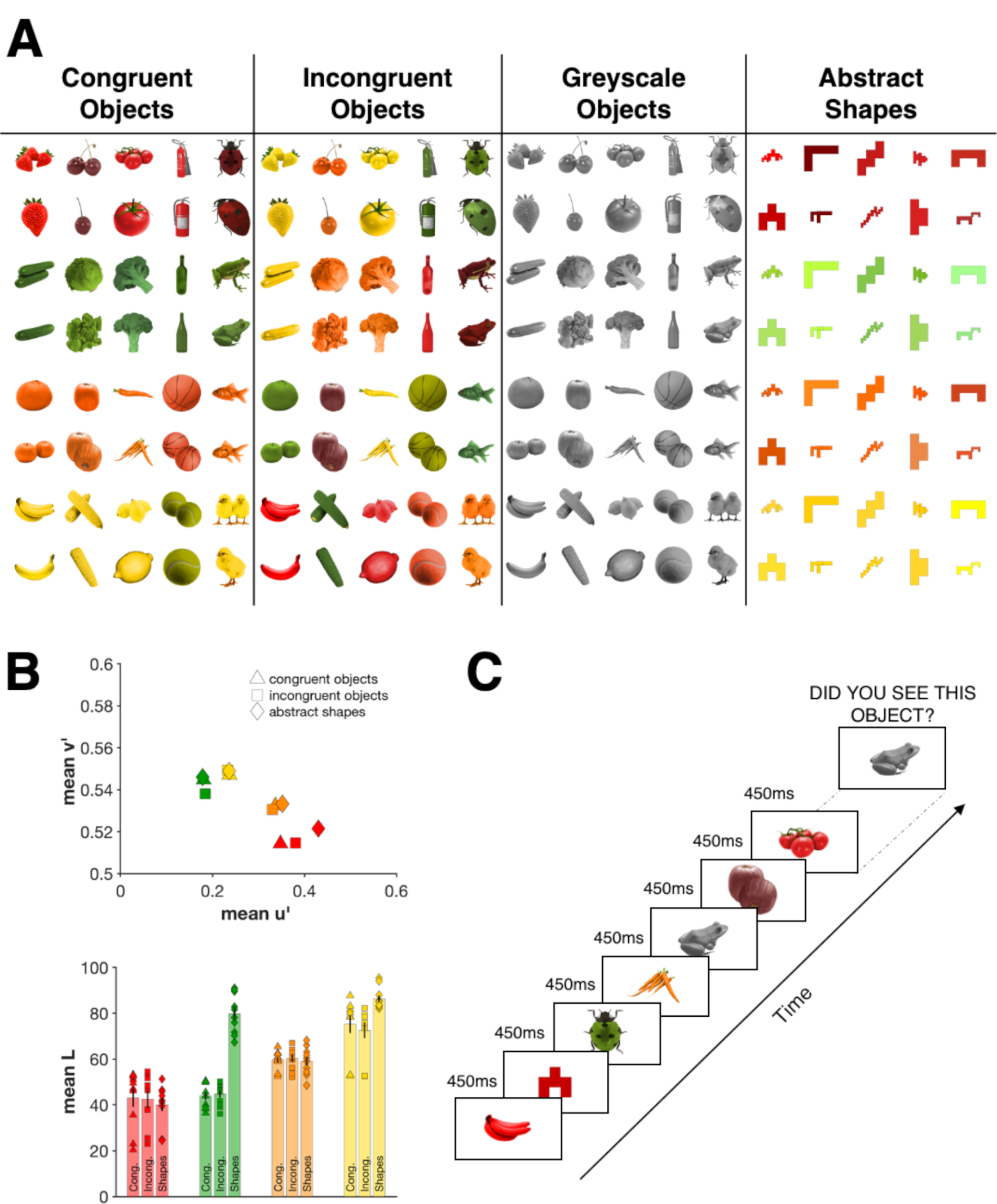
(A) shows all stimuli used in this experiment. The same objects were used in the congruent, incongruent, and greyscale conditions. There were two exemplars of each object. Colours in the congruent and incongruent condition were matched. The abstract shapes were identical across colour categories. (B) shows the mean chromaticity coordinates for the 2° observer under D65 illumination for each colour category (top) as well as the mean lightness of all coloured stimuli used in this experiment (bottom). The colours were transformed into CIELUV space using the OptProp toolbox (Wagberg, 2007). (C) shows an example sequence of the main task. Participants viewed each object for 450ms. After each sequence, one object was displayed and participants had to indicate whether they had seen this object in the previous sequence or not.

This procedure resulted in 40 congruent objects (10 of each colour), and 40 incongruent objects (10 of each colour, Figure 1). We added two additional stimulus types to this set: the full set of 40 greyscale images, and a set of 10 different angular abstract shapes, coloured in each of the four hues for a set of 40 (see Figure 1). As is clear in Figure 1, the colours of the abstract shapes appeared brighter than the colours of the objects, this is because the latter were made by overlaying hue on greyscale, whereas the shapes were simply coloured. As our principle goal was to ensure that the congruent objects appeared to have their typical colouring, we did not match the overall luminance of the coloured stimuli. For example, if we equated the red of a cherry with the yellow of a lemon, neither object would look typically coloured. Thus, each specific colour pair is not equated for luminance; however, we have the same colours across different conditions.

All stimuli were presented at a distance of 114cm. To add visual variability, which reduces the low-level featural overlap between the images, we varied the image size from trial to trial by 2 degrees of visual angle. The range of visual angles was therefore between ∼4.3 – 6.3 degrees.

### Experimental Design and Statistical Analysis

#### Experimental tasks

In the main task (Figure 1C), participants completed eight blocks of 800 stimulus presentations each. Each individual stimulus appeared 40 times over the course of the experiment. Each stimulus was presented centrally for 450ms with a black fixation dot on top of it. To keep participants attentive, after every 80 stimulus presentations, a target image was presented until a response was given indicating whether this stimulus had appeared in the last 80 stimulus presentations or not (50% present vs absent). The different conditions (congruent, incongruent, grey-scale, abstract shape) were randomly intermingled throughout each block, and the target was randomly selected each time. On average, participants performed with 90% (SD=5.4%) accuracy.

After completing the main blocks, we collected behavioural object-naming data to test for a behavioural congruency effect with our stimuli. On the screen, participants saw each of the objects again (congruent, incongruent or greyscale) in a random order and were asked to name the objects as quickly as possible. As soon as voice onset was detected, the stimulus disappeared. We marked stimulus-presentation times with a photodiode and recorded voice-onset with a microphone. Seventeen participants completed three blocks of this reaction time task, one participant completed two blocks, and for two participants we could not record any reaction times. Each block contained all congruent, incongruent and grey-scale objects presented once.

Naming reaction times were defined as the difference between stimulus-onset and voice-onset. Trials containing naming errors and microphone errors were not analysed. We calculated the median naming time for each exemplar for each person and then averaged the naming times for each of the congruent, incongruent and greyscale conditions.

#### MEG data acquisition

While participants completed the main task of the experiment, neuromagnetic recordings were conducted with a whole-head axial gradiometer MEG (KIT, Kanazawa, Japan), containing 160 axial gradiometers. We recorded the MEG signal with a 1000Hz frequency. An online low-pass filter of 200Hz and a high-pass filter of 0.03Hz were used. All stimuli were projected on a translucent screen mounted on the ceiling of the magnetically shielded room. Stimuli were presented using MATLAB with Psychtoolbox extension (Brainard, 1997; Brainard & Pelli, 1997; Kleiner et al., 2007). Parallel port triggers and the signal of a photodiode were used to mark the beginning and end of each trial. A Bimanual 4-Button Fiber Optic Response Pad (Current Designs, Philadelphia, USA) was used to record the responses.

Before entering the magnetically shielded room for MEG recordings, an elastic cap with five marker coils was placed on the participant’s head. We recorded head shape with a Polhemus Fastrak digitiser pen (Colchester, USA) and used the marker coils to measure the head position within the magnetically shielded room at the start of the experiment, halfway through and at the end.

#### MEG data analysis: Preprocessing

FieldTrip (Oostenveld et al., 2011) was used to preprocess the MEG data. The data were downsampled to 200Hz and then epoched from -100 to 500ms relative to stimulus onset. We did not conduct any further preprocessing steps (filtering, channel selection, trial-averaging etc.) to keep the data in its rawest possible form.

#### MEG data analysis: Decoding Analyses

For all our decoding analyses, patterns of brain activity were extracted across all 160 MEG sensors at every timepoint, for each participant separately. We used a regularised linear discriminant analysis (LDA) classifier which was trained to distinguish the conditions of interest across the 160-dimensional space. We then used independent test data to assess whether the classifier could predict the condition above chance in the new data. We conducted training and testing at every timepoint and tested for significance using random-effects Monte Carlo cluster (TFCE; Smith & Nichols, 2009) statistics, corrected for multiple comparisons using the max statistic across time points (Maris & Oostenveld, 2007). Note that our aim was not to achieve the highest possible decoding accuracy, but rather to test whether the classifier could predict the conditions above chance at any of the timepoints (i.e., “classification for interpretation”, Hebart & Baker, 2017). Therefore, we followed a minimal preprocessing pipeline and performed our analyses on a single-trial basis. Classification accuracy above chance indicates that the MEG data contains information that is different for the categories. We used the CoSMoMVPA toolbox (Oosterhof et al., 2016) to conduct all our analyses.

We ran several decoding analyses which can be divided in three broad themes. First, we tested when we can differentiate between trials where congruently and incongruently coloured objects were presented. This gives us an indication of the timecourse of the integration of visual object representations and stored conceptual knowledge. Second, we examined single feature processing focusing on colour processing and how the typicality of object-colour combinations influences colour processing over time. Third, we looked at another single feature, shape, and tested whether object-colour combinations influence shape processing over time.

For the congruency analysis (Figure 2A), we tested whether activation patterns evoked by congruently coloured objects (e.g., red strawberry) differ from activation patterns evoked by incongruently coloured objects (e.g., yellow strawberry). Any differential response that depends on whether a colour is typical or atypical for an object (a congruency effect) requires the perceived shape and colour to be bound and compared to a conceptual object representation activated from memory. We trained the classifier on all congruent and incongruent trials *except* for trials corresponding to one pair of matched exemplars (e.g., all instances of congruent and incongruent strawberries and congruent and incongruent bananas). We then tested the classifier using only the left-out exemplar pairs. We repeated this process until each matched exemplar pair had been left out (i.e., used as test data) once. Leaving an exemplar pair out ensures that there are identical objects and colours for both classes (congruent and incongruent) in both the training and the testing set, and that the stimuli of the test set have different shape characteristics than any of the training objects. As such, the only distinguishing feature between the conditions is the *conjunction* of shape and colour features, which defines congruency. This allows us to compare directly whether (and at which timepoint) stored object representations interacts with incoming object-colour information.

**Figure 2.**
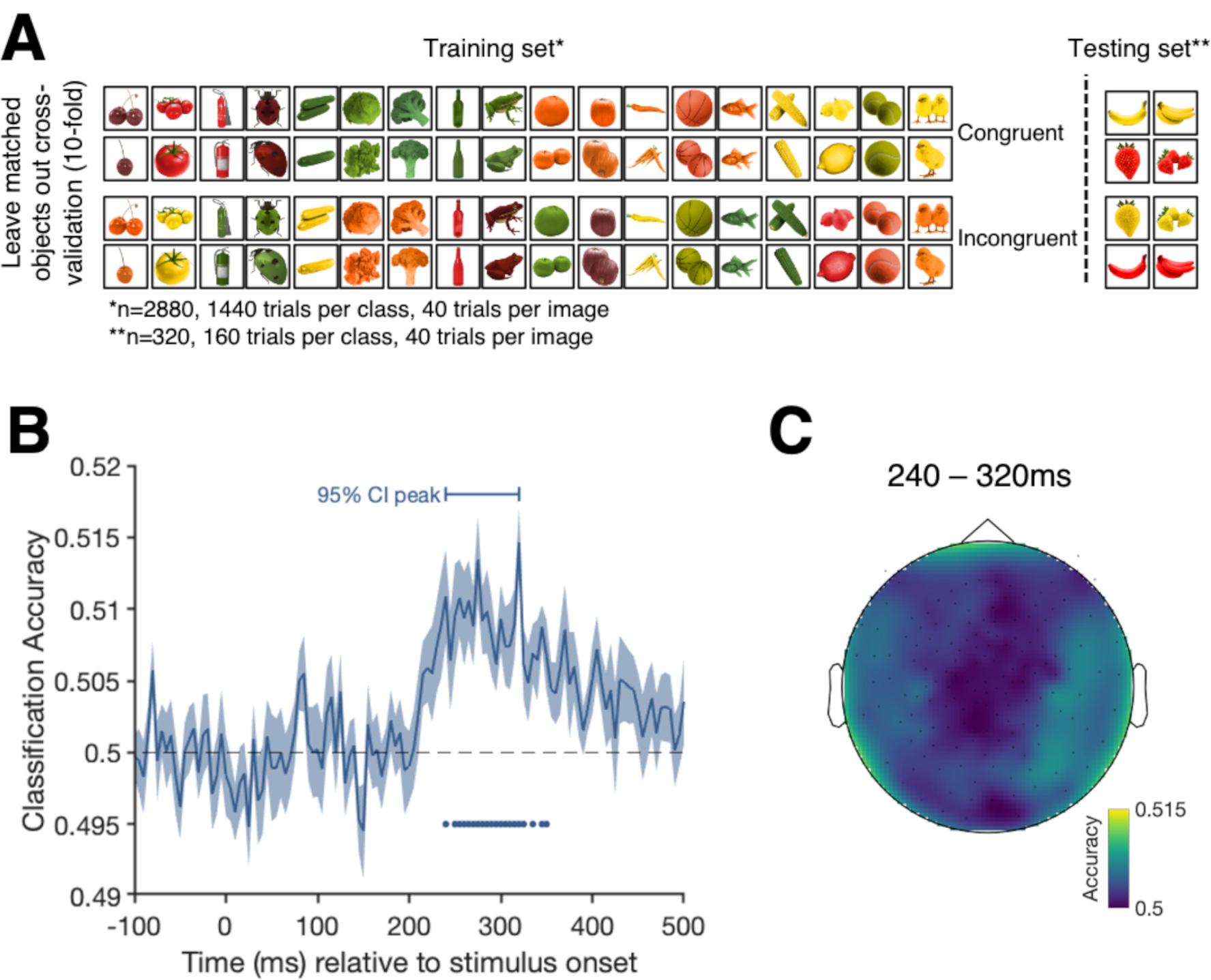
Cross-validation and results of the congruency analysis contrasting trials from the congruent and incongruent conditions. (A) shows the leave-one-matched-exemplar-out cross validation approach for a single fold for the congruency decoding analysis. The classifier was trained on the trials shown on the left and tested on the trials on the right, ensuring that the classifier is not tested on the exemplars that it trained on. This limits the effect features other than congruency can have on classifier performance. (B) shows the classification accuracy over time. Shading represents the standard error across participants. Black dashed line represents chance level (50% - congruent versus incongruent). Filled dots highlight significant timepoints, corrected for multiple comparisons. The horizontal bar above the curve shows the 95% confidence interval of the peak. (C) is an exploratory sensor searchlight analysis in which we run the same analysis across small clusters of sensors. The colours highlight the decoding accuracy for each sensor cluster averaged over the 95% confidence interval of the peak timepoints.

Next, we focused on the timecourse of colour processing. First, we examined the timecourse of colour processing independent of congruency (Figure 3A). For this analysis, we trained the classifier on distinguishing between the four different colour categories of the abstract shapes and tested its performance on an independent set of abstract shape trials. We always left one block out for the cross-validation (8-folds). The results of this analysis give an indication about the emergence of when the representations differ between different surface colours, but as we did not control the colours to be equal in luminance or have the same hue difference between each pair, this is not a pure chromatic measure. We did not control luminance because we used these colours to create our coloured objects, which needed to look as realistic as possible. Thus, the colour decoding analysis includes large and small differences in hue *and in luminance* between the categories. To look at the differences between each colour pair, we also present confusion matrices showing the frequencies of the predicted colour categories at peak decoding.

**Figure 3.**
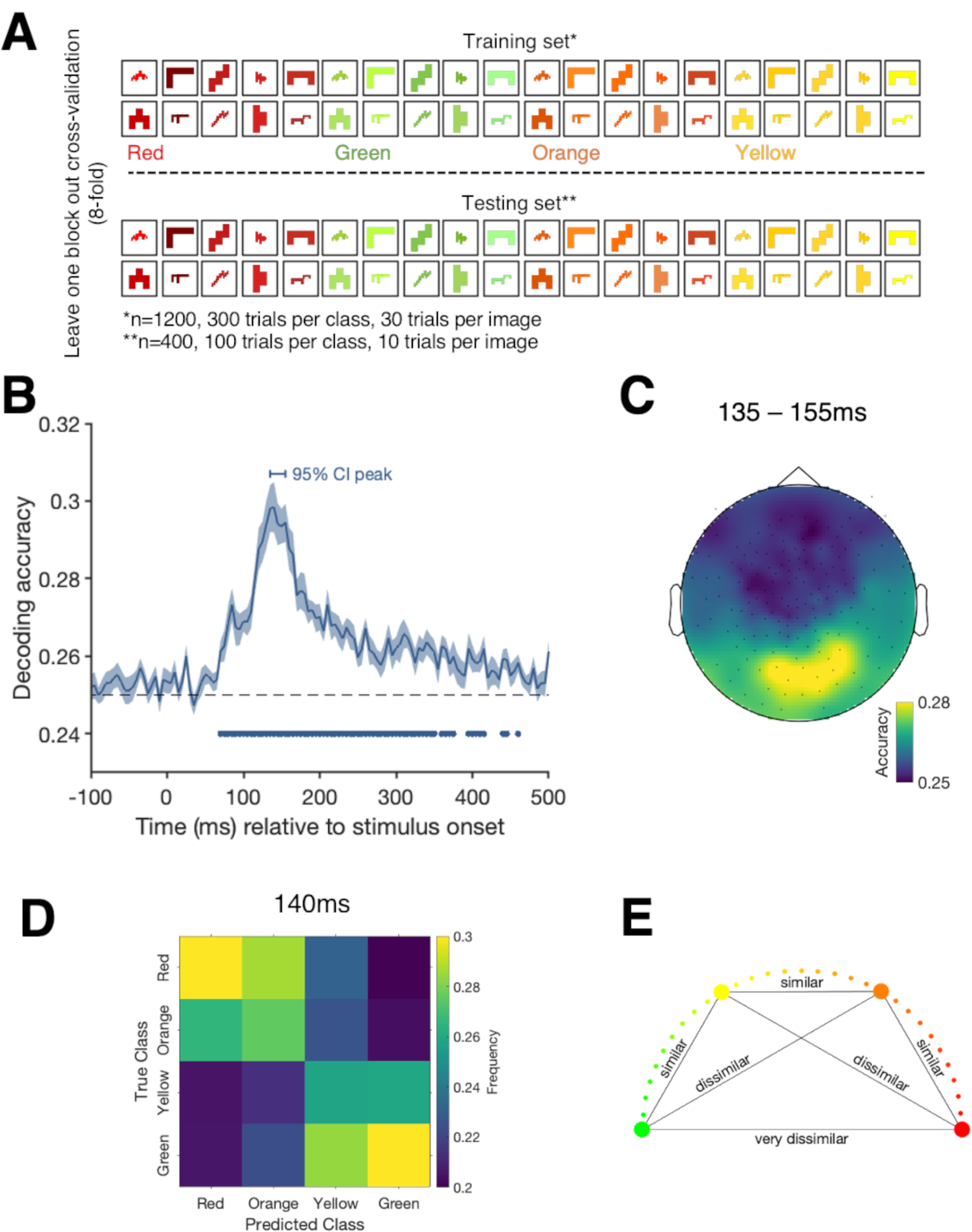
(A) depicts the colour decoding analysis when training the classifier to distinguish between the different colour categories of the abstract shapes and testing on a block of independent abstract shape trials. (B) shows the decoding accuracy for the colour decoding analysis over time. Shading represents the standard error across participants. Black dashed line represents chance level (25% - red versus green versus orange versus yellow). Filled dots highlight significant timepoints, corrected for multiple comparisons. The horizontal bar above the curve shows the 95% confidence interval of the peak. (C) shows the results of a exploratory searchlight analysis over small sensor clusters averaged across the timepoints of the 95% confidence interval for peak decoding. Colours indicate the decoding accuracies at each sensor. (D) depicts a confusion matrix for peak decoding (140ms) showing the frequencies at which colour categories were predicted given the true class. (E) shows the similarity of the colour categories which might underlie the results in (D).

Our second colour processing analysis was to examine whether the conjunction of object and colour influenced colour processing (Figure 4A). Perceiving a strongly associated object in the context of viewing a certain colour might lead to a more stable representation of that colour in the MEG signal. For example, if we see a yellow banana, the banana shape may facilitate a representation of the colour yellow earlier than if we see a yellow strawberry. To assess this possibility, we trained the classifier to distinguish between the surface colours of the abstract shapes (i.e., red, orange, yellow, green; chance: 25%). We then tested how well the classifier could predict the colour of the congruent and incongruent objects. Training the classifier on the same abstract shapes across colour categories makes it impossible that a certain shape-colour combination drives an effect, as the only distinguishing feature between the abstract shapes is colour. This analysis allows us to compare whether the typicality of colour-form combinations has an effect on colour processing.

**Figure 4.**
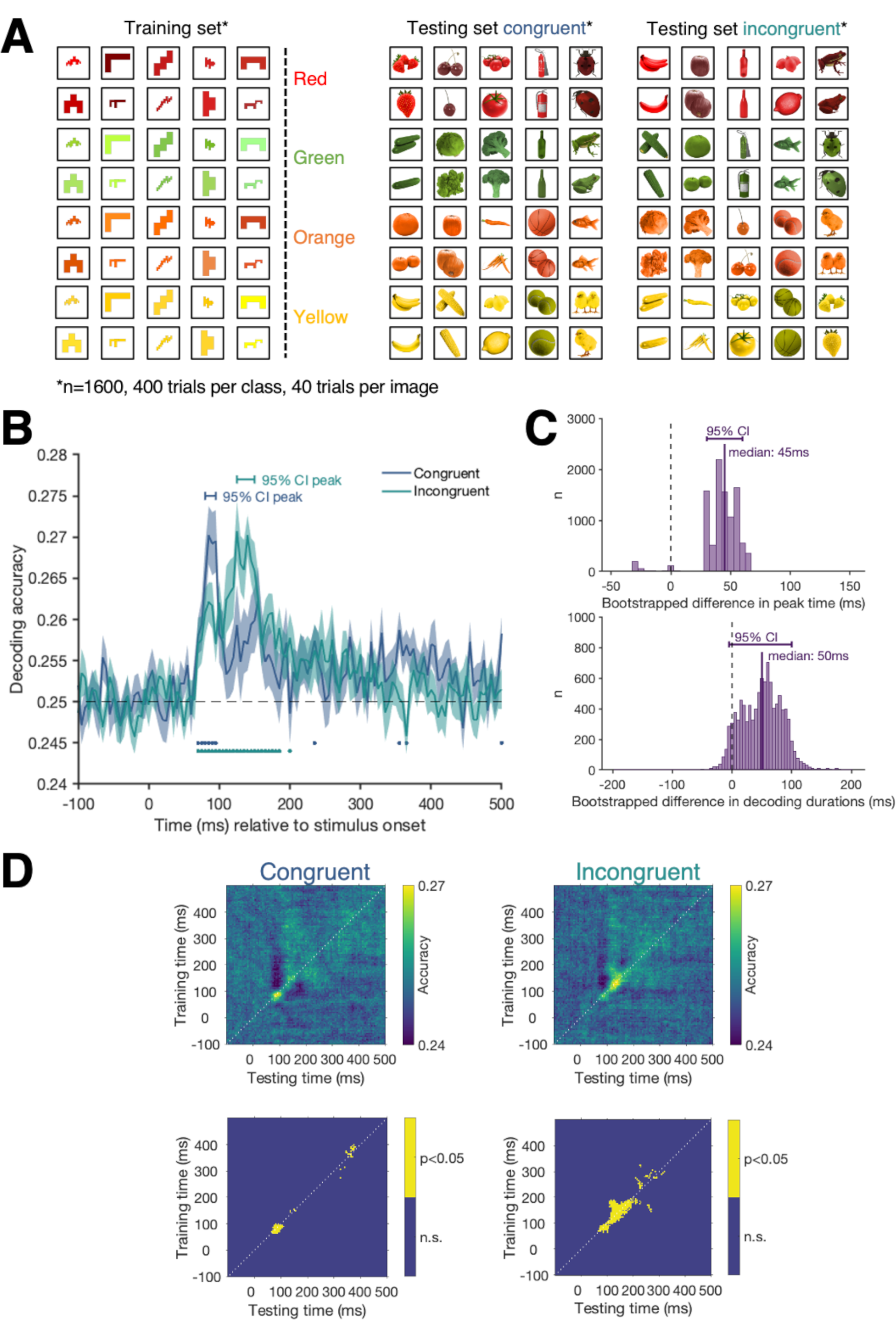
Results of the colour decoding analysis for the congruent and incongruent trials. Here, the classifier was trained to distinguish the colour of all abstract shape trials and tested on the congruent and incongruent trials separately (A). (B) shows the classification accuracy over time for the congruent (blue) and incongruent (green) trials. Shading represents the standard error across participants. Black dashed line indicates chance level (25% - red versus green versus orange versus yellow). Blue (congruent) and green (incongruent) dots highlight timepoints at which we can decode the surface colour significantly above chance, corrected for multiple comparisons. (C) shows the bootstrapped differences in peak time (top) and the bootstrapped differences in decoding duration (bottom) for the congruent and the incongruent conditions. (D) shows the results of the same analysis across all possible training and testing timepoint combinations. These time-time matrices allow us to examine how the signal for the congruent colours (left) and incongruent colours (right) evolves over time. The top row shows the classification accuracy at every timepoint combination with lighter pixels reflecting higher decoding accuracies. The bottom row shows clusters where decoding is significantly above chance (yellow), corrected for multiple comparisons.

In our final set of analyses, we examined the timecourse of shape processing. First, to assess the timecourse of shape processing independent of congruency, we trained a classifier to distinguish the five different abstract shapes in a pairwise fashion (Figure 5A). We always used one independent block of abstract shape trials to test the classifier performance (8-fold cross-validation). The results of this analysis indicate when information about different shapes are is present in the neural signal, independent of other object features (e.g., colour) or congruency. Second, we tested whether the conjunction of object and colour has an effect on object decoding (Figure 6A). If object-colour influences early perceptual processes, we might see a facilitation for decoding objects when they are coloured congruently or interference when the objects are coloured incongruently. We used the greyscale object trials to train the classifier to distinguish between all of the objects. The stimulus set contained two exemplars of each item (e.g., strawberry 1 and strawberry 2). We used different exemplars for the training and testing set to minimise the effects of low-level visual features, however, given that there are major differences in object shapes and edges, we can still expect to see strong differences between the objects. The classifier was trained on one exemplar of all of the greyscale trials. We then tested the classifier’s performance on the congruent and incongruent object trials using the exemplars the classifier did not train on. We then swapped the exemplars used for training and testing set until every combination had been used in the testing set. Essentially, this classifier is trained to predict which object was presented to the participant (e.g., was it a strawberry or a frog?) and we are testing whether there is a difference depending on whether the object is congruently or incongruently coloured.

**Figure 5.**
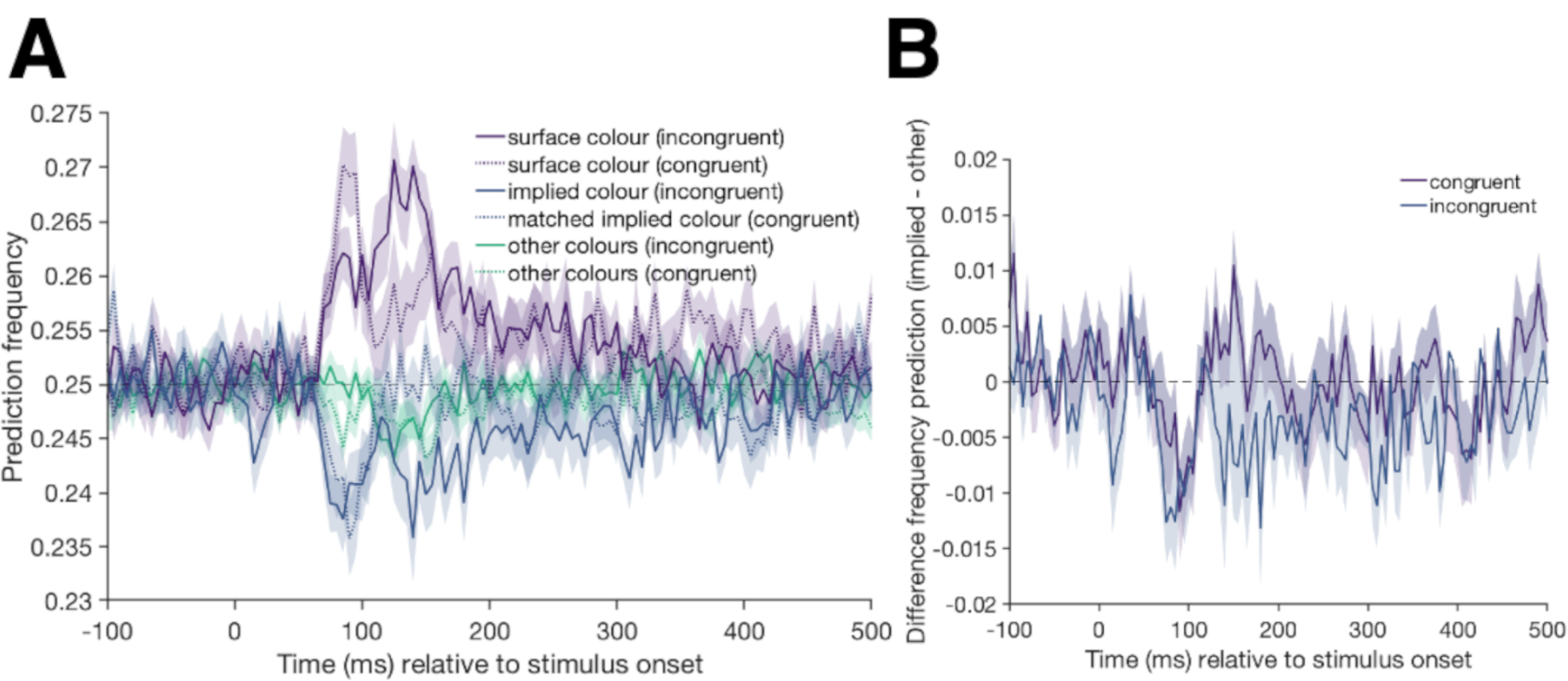
(A) shows the frequency of a predicted class when the classifier is trained on distinguishing colours in the abstract shape condition and tested on trials from the congruent (dotted lines) and incongruent (full lines) conditions. Shading represents the standard error across participants. There are clear peaks for the correct prediction of the surface colour between 100and 150ms (purple lines). In cases where the classifier makes an error, there is no evidence that the classifier picks the implied object colour (blue lines) more frequently than the other incorrect colours (green lines). Note that the classifier is trained on the abstract shape condition which has an uneven colour similarity, the errors in the incongruent condition have to be interpreted in relation to how often the matched implied colour in the congruent condition is predicted. (B) shows the difference of the classifier predicting the implied over the other colours for the congruent (purple) and incongruent (blue) conditions.

**Figure 6.**
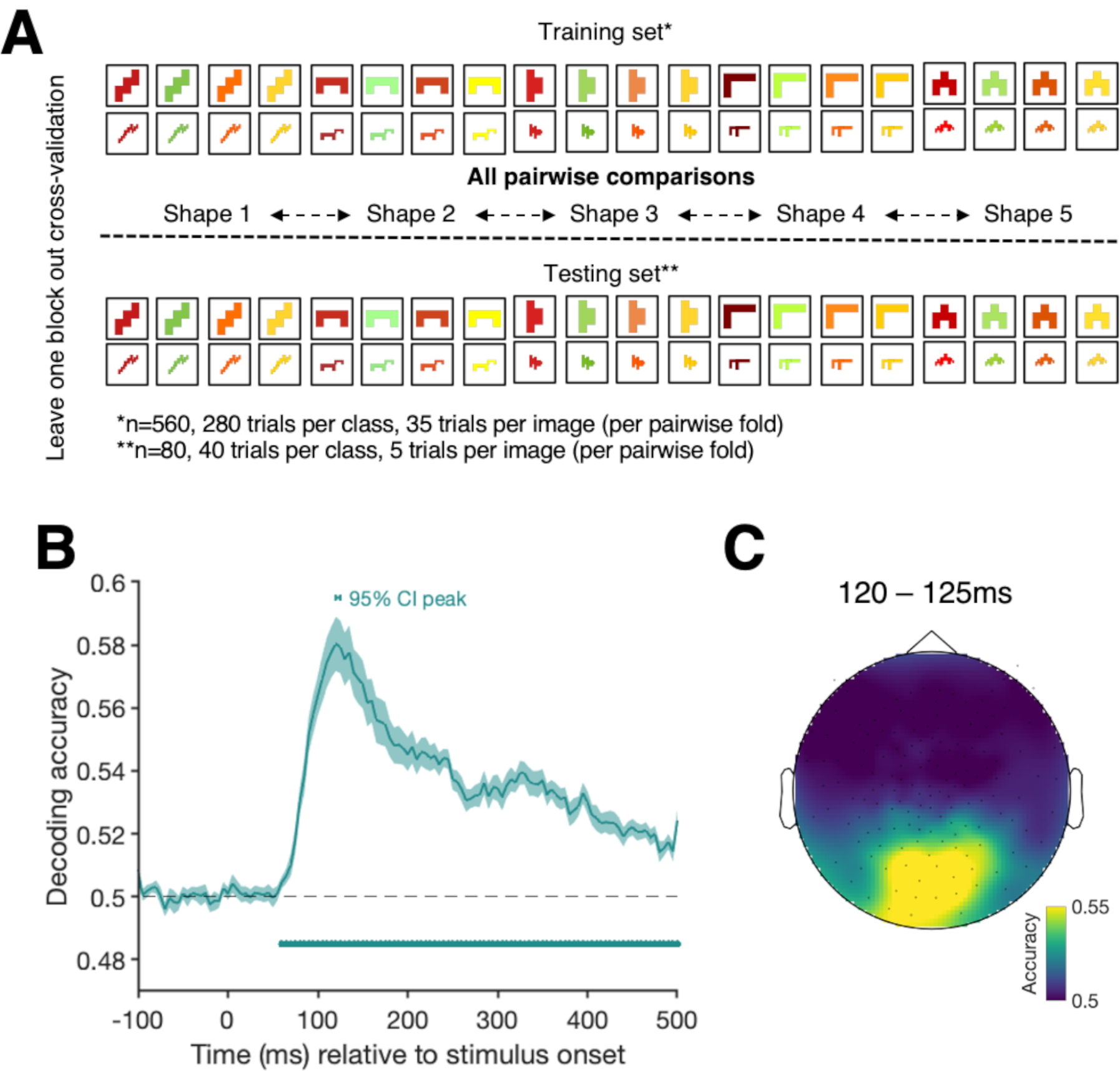
(A) depicts the shape decoding analysis when training the classifier to distinguish between the different categories of the abstract shapes and testing on a block of independent abstract shape trials. (B) shows the decoding accuracy for the shape decoding analysis over time. Shading represents the standard error across participants. Black dashed line represents chance level (50% - pairwise comparison of all shapes). Filled dots highlight significant timepoints, corrected for multiple comparisons. The horizontal bar above the curve shows the 95% confidence interval of the peak. (C) shows the results of an exploratory searchlight analysis over small sensor clusters averaged across the timepoints of the 95% confidence interval for peak decoding. Colours indicate the decoding accuracies at each sensor.

#### Statistical Inferences

In all our analyses, we used random effects Monte-Carlo cluster statistic using Threshold Free Cluster Enhancement (TFCE, Smith & Nichols, 2009) as implemented in the CoSMoMVPA toolbox to see whether the classifier could predict the condition of interest above chance. The TFCE statistic represents the support from neighbouring time points, thus allowing for detection of sharp peaks and sustained small effects over time. We used a permutation test, swapping labels of complete trials, and re-ran the decoding analysis on the data with the shuffled labels 100 times per participant to create subject-level null-distributions. We then used Monte-Carlo sampling to create a group-level null-distribution consisting of 10,000 shuffled label permutations for the time-resolved decoding, and 1000 for the time-generalisation analyses (to reduce computation time). The null distributions were then transformed into TFCE statistics. To correct for multiple comparisons, the *maximum* TFCE values across time in each of the null distributions was selected. We then transformed the true decoding values to TFCE statistics. To assess whether the true TFCE value at each timepoint is significantly above chance, we compared it to the 95^th^ percentile of the corrected null distribution. Selecting the maximum TFCE value provides a conservative threshold for determining whether the observed decoding accuracy is above chance, corrected for multiple comparisons.

To assess at which timepoint the decoding accuracy peaks, we bootstrapped the participants’ decoding accuracies for each analysis 10,000 times and generated 95% confidence intervals for peak decoding. For the analyses in which we are comparing colour and exemplar decoding for congruent and incongruent trials, we also compared the above chance decoding durations.

To test for the duration of above chance decoding, we bootstrapped the data (10,000 times) and ran our statistics. At each iteration we then looked for the longest period in which we have above chance decoding in consecutive timepoints. We plotted the bootstrapped decoding durations and calculated medians to compare the distributions for the congruent and the incongruent condition.

## Results

### Behavioural results

We first present the data from our behavioural object-naming task to confirm that our stimuli induce a congruency effect on object naming times. All incorrect responses and microphone errors were excluded from the analysis (on average across participants: 10.1%). We then calculated the median reaction time for naming each stimulus. If a participant named a specific stimulus incorrectly across trials (e.g., incongruently coloured strawberry was always named incorrectly), we removed this stimulus completely to ensure that the reaction times in one condition were not skewed. We ran a repeated measures ANOVA to compare the naming times for the different conditions in the behavioural object naming task using JASP (Love et al., 2015). Naming times were significantly different between the conditions (F(2,34) = 12.8; p<0.001). Bonferroni-corrected post hoc comparisons show that participants were faster to name the congruently coloured (701ms) than the incongruently coloured (750ms) objects (p_bonf_ < 0.001; 95%CI for mean difference [23.8, 72.8]). It took participants on average 717ms to name the greyscale objects which was significantly faster than naming the incongruently coloured objects (p_bonf_ = 0.007; 95%CI for mean difference [7.8, 56.8]) but not significantly slower than naming the congruently coloured objects (p_bonf_ = 0.33.; 95%CI for mean difference [-40.5, 8.5]). These results suggest that the objects we used here do indeed have associations with specific canonical colours, and we replicate that these objects are consistently associated with a particular colour (Bannert & Bartels, 2013; Joseph, 1997; Lloyd-Jones et al., 2012; Naor-Raz et al., 2003; Tanaka & Presnell, 1999; Therriault et al., 2009).

In the main task, participants were asked to indicate every 80 trials whether they had seen a certain target object or not. The aim of this task was to keep participants motivated and attentive throughout the training session. On average, participants reported whether the targets were present or absent with 90% accuracy (SD = 5%, range: 81.25% - 100%).

### MEG results

The aim of our decoding analyses was to examine the interaction between object-colour knowledge and object representations. First, we tested for a difference in the brain activation pattern for congruently and incongruently coloured objects. The results show distinct patterns of neural activity for congruent compared to incongruent objects in a cluster of consecutive timepoints stretching from 250 to 325ms after stimulus onset, demonstrating that brain activity is modulated by colour congruency in this time window (Figure 2B). Thus, binding of colour and form must have occurred by ∼250ms and stored object-colour knowledge is integrated with incoming information. An exploratory searchlight (Carlson et al., 2019; Collins et al., 2018; Kaiser et al., 2016) across small clusters (9 at a time) of MEG sensors suggests that this effect is driven a range of frontal, temporal and parietal sensor clusters (Figure 2C).

To examine the timecourse of colour processing separately from congruency, we decoded the surface colours of the abstract shapes (Figure 3A). Consistent with earlier results (Teichmann et al., 2019), we found that colour can be decoded above chance from the abstract shape trials in a cluster stretching from 70 to 350ms (Figure 3B). Looking at data from an exploratory sensor searchlight analysis across small clusters of sensors shows that colour information at peak decoding is mainly distinguishable from occipital and parietal sensors. To examine whether all colours could be dissociated equally well, we also looked at confusion matrices displaying how frequently each colour category was predicted for each colour (Figure 3D). The results show that at the decoding peak (140ms), red and green are most easily distinguishable and that the prediction errors are not equally distributed: Red trials are more frequently misclassified as being orange than green or yellow and green trials are more frequently misclassified as being yellow than orange or red. This indicates that colours that are more similar evoke a more similar pattern of activation than colours that are dissimilar (Figure 3E).

To assess whether congruency influences colour processing, we trained a classifier to distinguish between the colours in the abstract shape condition and then tested it on the congruent and incongruent trials separately (Figure 4A). Colour can be successfully classified in a cluster stretching from 75 to 125ms for the congruent condition and in a cluster stretching from 75 to 185ms for the incongruent trials (Figure 4B). These results suggest there may be a difference in the way colour information is processed depending on the congruency of the image, specifically evident in the decoding peaks and decoding duration. To test whether there is a true difference in decoding timecourses, we bootstrapped the data and looked at the peak decoding and the longest consecutive streak of above chance decoding. Comparing the peak decoding times for the congruent and the incongruent condition, we find that they are different from each other (Figure 4C, top). However, comparing the confidence intervals of the decoding durations we find no consistent differences between the congruent and the incongruent condition (Figure 4C, bottom). This could be due to the fact that on- and offsets in above chance decoding are affected by signal strength and thresholds (cf. Grootswagers et al., 2017). The peak differences are a more robust measure and suggest that colour stronger colour decoding occurs later in the incongruent compared to congruent condition. To get a complete picture of how these signals evolve over time, we used time-generalisation matrices (Figure 4D and 4E). To create time-generalisation matrices, we trained the classifier on each timepoint of the training dataset and then tested it on all timepoints of the test set. The diagonal of these matrices corresponds to the standard time-resolved decoding results (e.g., training at 100ms and testing at 100ms). A decodable off-the-diagonal effect reflects a temporal asynchrony in information processing in the training and testing set (cf. Carlson et al., 2011; King & Dehaene, 2014). Our data show that colour category was decodable from both conditions early on (∼70ms). In the incongruent condition, the activation associated with colour seems to be sustained longer (Figure 4E) than for the congruent condition (Figure 4D), but for both, decoding above chance occurs mainly along the diagonal. This suggests that the initial pattern of activation for colour signals occurs at the same time but that the signals associated with colour are prolonged when object-colour combinations are unusual relative to when they are typical.

In an exploratory colour analysis, we also examined which errors the classifier made when predicting the colour of the incongruently coloured objects. We looked at whether the implied object colour is predicted more often than the surface colour or the other colours. However, as errors were not equally distributed across the incorrect labels in the training (abstract shape) dataset, we need to compare the misclassification results for the incongruent condition to the results from the congruent condition, to take these differing base rates into account. For each object in the incongruent condition (e.g., yellow strawberry), we have a colour-matched object in the congruent condition (e.g., yellow banana). We made use of these matched stimuli by looking at misclassifications and checking how frequently the implied colour of an incongruent object (e.g., red for a yellow strawberry) was predicted in comparison to the matched congruent object (e.g., red for a yellow banana). If the implied colour of incongruently coloured objects was activated along with the surface colour, we should see a higher rate of implied colour predictions (e.g., red) for the incongruent object (e.g., yellow strawberry) than for the colour-matched congruent object (e.g., yellow banana).

The results (Figure 5) do not show this pattern: at the first peak (∼80-110ms), the “other” colours are actually more likely to be chosen by the classifier than the implied colour, for both the congruent and incongruent condition. A possible explanation for not seeing an effect of implied colour in the colour decoding analysis is that the decoding model is based on the actual colour pattern, whereas the timing and mechanisms of implied colour activation may be different (Teichmann et al., 2019).

The goal of the third analysis was to examine whether shape representations are affected by colour congruency. It could be the case, for example, that the representation of banana-shapes compared to strawberry-shapes is enhanced when their colours are correct. First, we tested when shape representations can be decoded independent of colour congruency. We trained the classifier to distinguish between the five different abstract shapes in a pairwise fashion and then tested its performance on independent data (Figure 6A). The data show that shape information can be decoded in a cluster stretching from 60 to 500ms (Figure 6B). Running an exploratory searchlight analysis on small clusters of sensors (9 at a time) shows that shape information at peak decoding is mainly driven by occipital sensors (Figure 6C).

To examine whether colour affects object processing, we trained a classifier to distinguish between trials in which the participant saw one of the exemplars of each of the twenty objects in greyscale (e.g., greyscale strawberry 1, greyscale cherry 1, etc.). We then tested at which timepoint the classifier could successfully cross-generalise to the other exemplar of that object in the congruent and incongruent condition separately (Figure 7A). If object representations (e.g., banana) are influenced by the typicality of their colours, then cross-generalisation should be different for congruent and incongruent trials. Note that although the exact images are unique, there are shared shape characteristics between exemplars (e.g., the two frog exemplars share some shape aspects despite being different postures), which can be expected to drive an effect. The results show the neural data has differential information about the object in a cluster stretching from 65 to 500ms for both the congruent and the incongruent test sets (Figure 7B). These results show that we can decode the object class early on, at a similar time to when we can decode the abstract shape conditions, suggesting that the classifier here is driven strongly by low-level features (like shape), rather than being influenced by colour congruency. The timecourse for congruent and incongruent object decoding is very similar in terms of peak decoding and decoding duration (Figure 7C). Thus, our data suggest that there is no effect of colour congruency on object processing.

**Figure 7.**
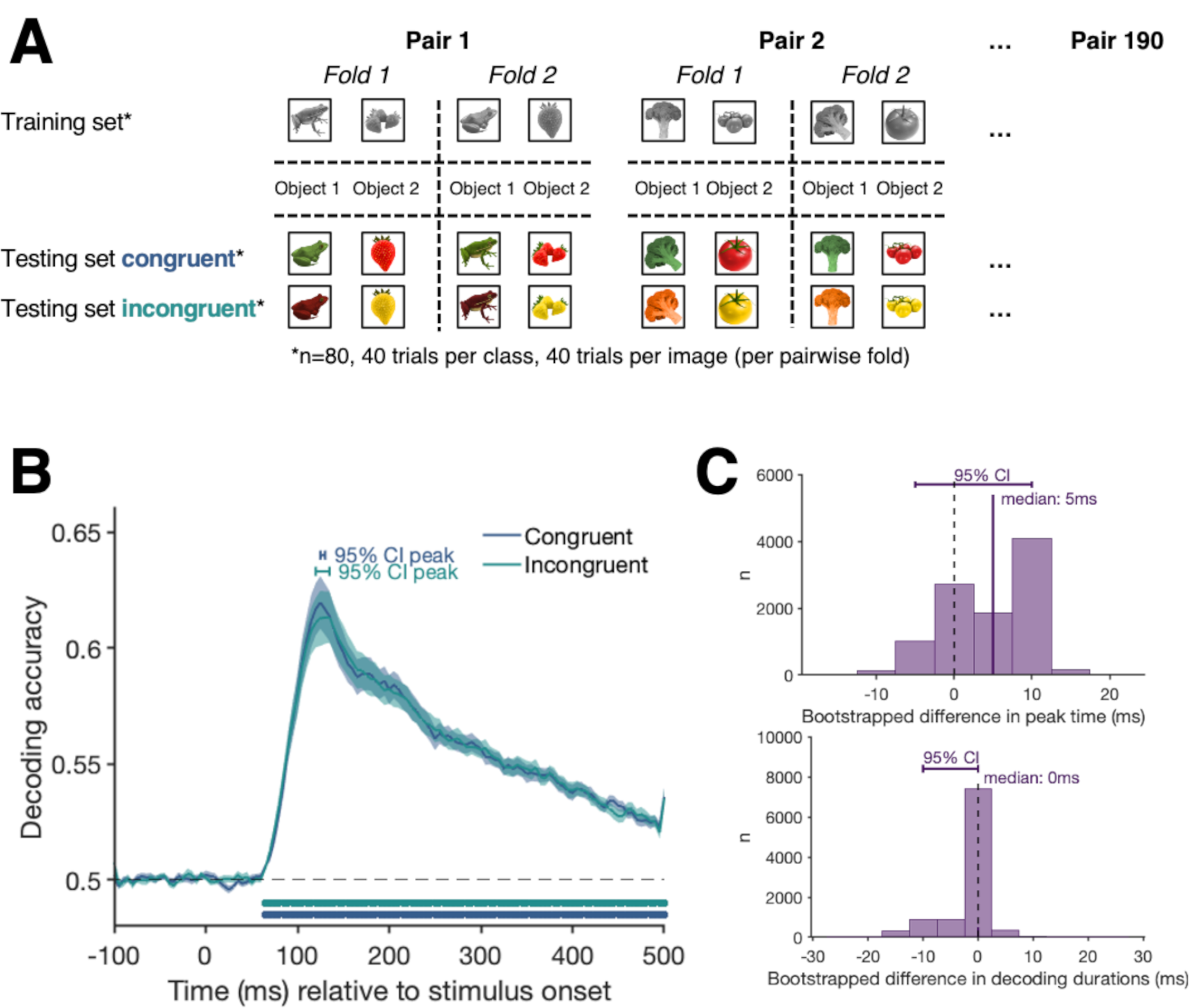
Results of the object exemplar decoding analysis. The classifier was trained to distinguish between all pairwise object categories in the greyscale object condition. We used one exemplar of each class for the training and the other exemplar for testing the classifier. Testing was done for the congruent and incongruent trials separately (A). (B) shows the classification accuracy over time for the object decoding analysis when testing the classifier’s performance on congruent (blue) and incongruent (green) trials. Shading represents the standard error across participants. Black dashed line represents chance level (50% - pairwise decoding for all 20 different object categories). Blue (congruent) and green (incongruent) dots highlight significant timepoints (p<0.05), corrected for multiple comparisons. (C) shows the bootstrapped differences in peak time (top) and the bootstrapped differences in decoding duration (bottom) for the congruent and the incongruent conditions.

Overall, the results here show that single features present within the incoming visual stimuli are decodable earlier than the congruency between them, which can be seen as an index for accessing stored conceptual knowledge (Figure 8). When we compare colour and shape decoding for abstract shapes and for congruently and incongruently coloured objects, the decoding onsets are very similar, suggesting the initial processes of single feature processing are not influenced by congruency. However, peak colour decoding occurs later for incongruently coloured in comparison to congruently coloured objects suggesting that colour congruency influences colour processing to some degree.

**Figure 8.**
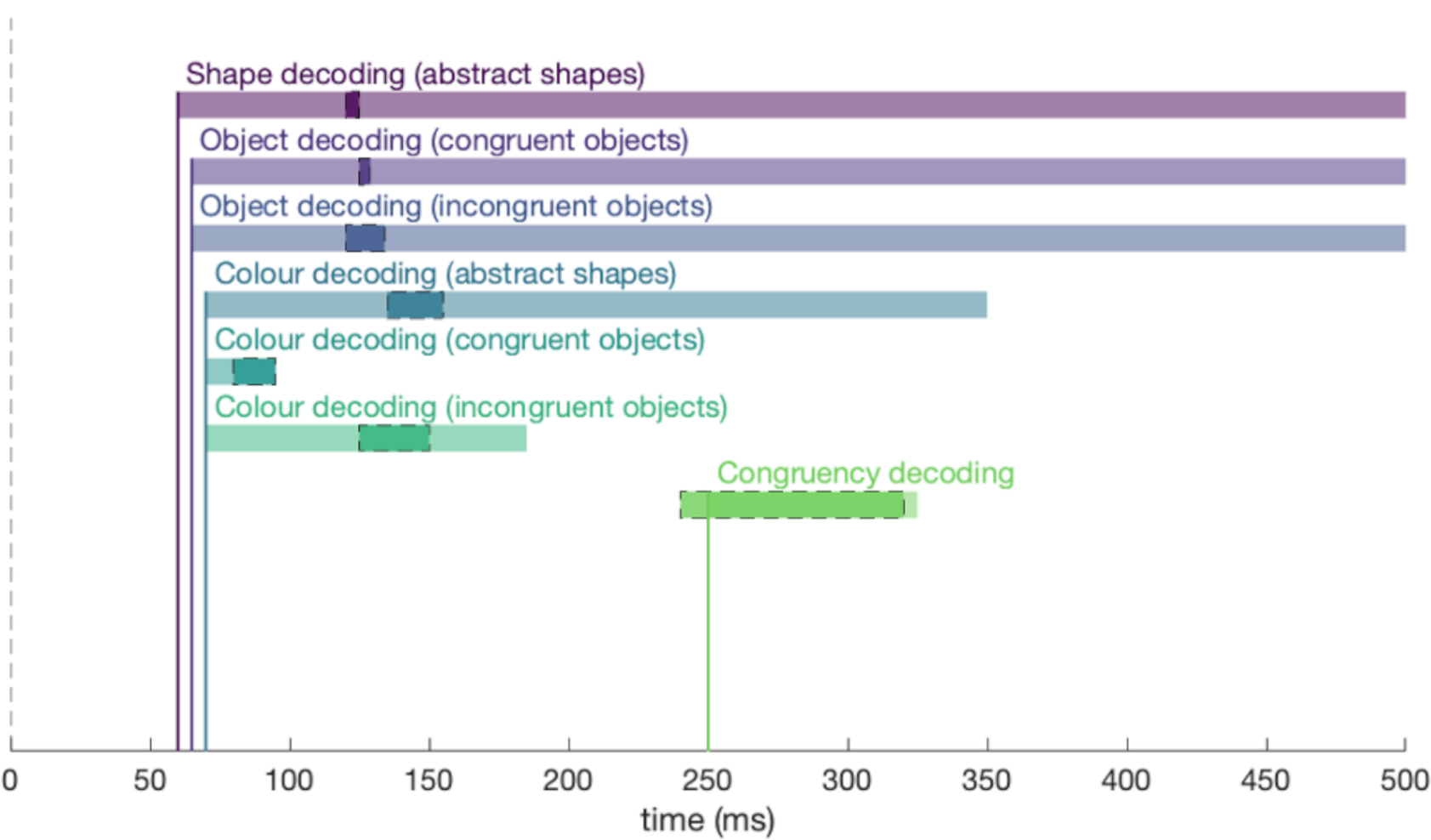
Overview of the findings. Each coloured bar shows the the onset (x axis) and duration (length of coloured bar) at which feature and conjunction information was contained in the neural signal. Darker shadings surrounded by dotted black lines show the bootstrapped 95% confidence interval for peak decoding. The dotted vertical line represents stimulus onset.

## Discussion

A crucial question in object recognition concerns how incoming visual information interacts with stored object concepts to create meaningful vision under varying situations. The aims of the current study were to examine the temporal dynamics of object-colour knowledge and to test whether activating object-colour knowledge influences the early stages of colour and object processing. Our data provide three major insights: First, congruently and incongruently coloured objects evoke a different neural representation after ∼250ms suggesting that, by this time, visual object features are bound into a coherent representation and compared to stored object representations. Second, colour can be decoded at a similar latency (∼70ms) irrespective of whether participants view coloured abstract shapes, or congruently and incongruently coloured objects. However, peak decoding occurs later when viewing incongruently coloured objects compared to congruent ones. Third, we do not find an influence of colour congruency on object processing, which may suggest that behavioural congruency effects are due to conflict at a later stage in processing.

Colour congruency can act as an index to assess when prior knowledge is integrated with bound object features. When comparing brain activation patterns of the same objects presented in different colours, there was a decodable difference between congruent and incongruent conditions from ∼250ms onwards suggesting a stored object representation that contains information about the typical colour of an object must have been activated by this stage. Prior to this time, the signal is primarily driven by processing of early perceptual features such as colour and shape, which were matched for the congruent and incongruent conditions (same objects, same colours, only the combination of colour and shape differed). Although from our data we cannot draw direct conclusions about which brain areas are involved in the integration of incoming visual information and stored object knowledge, our congruency analysis adds to the fMRI literature by showing the relative timecourse at which a meaningful object representation emerges. Activating object-colour knowledge from memory has been shown to involve the ATL (e.g., Coutanche & Thompson-Schill, 2014) and there is evidence that object-colour congruency coding occurs in perirhinal cortex (Price et al., 2017). Further support on the involvement of the ATL in the integration of incoming sensory information and stored representations comes from work on patients with semantic dementia (e.g., Bozeat et al., 2002) and studies on healthy participants using TMS (e.g., Chiou et al., 2014). Higher level brain areas in the temporal lobe have also been shown to be part of neuronal circuits involved in implicit imagery, supporting visual perception by augmenting incoming information with stored conceptual knowledge (e.g., Albright, 2012; Miyashita, 2004). The latency of congruency decoding here may thus reflect the time it takes to compare visual object representations with conceptual templates in higher-level brain areas such as the ATL, or the time it takes for feedback or error signals about colour congruency to arrive back in early visual areas.

Our results also show that colour congruency has an effect on colour processing. We found colour decoding onset at a similar time (∼70ms) for abstract shapes and congruently and incongruently coloured objects. This indicates that colour signals are activated initially independently of object shape, consistent with previous work showing that single features are processed first and that the conjunction of colour and shape occurs at a later stage (e.g., Seymour et al., 2015). However, we also found differences between colour processing in congruent and incongruent conditions: The colour signal peaked later in the incongruent relative to the congruent condition, suggesting that congruency influences the timecourse of colour processing to some degree. Our time-generalisation analysis (Figure 4D) supports this by showing that there is a different dynamic for congruent and incongruent trials. One possible explanation for this finding is that unusual feature pairings (e.g., shape and colour or texture and colour) might lead to local feedback signals that prolong colour processing. Alternatively, consistent with the memory colour literature (e.g., Hansen et al., 2006; Olkkonen et al., 2008; Witzel et al., 2011), it is possible that typical colours are co-activated along with other low-level features. For incongruent trials, this would then lead to two potential colours needing to be distinguished, extending the timeframe for processing and delaying the peak activation for the surface colour of the object.

The timecourse of exemplar decoding we present is consistent with previous studies on object recognition. Here, we found that exemplar identity could be decoded at ∼65ms. Similar latencies have been found in other M/EEG decoding studies (Carlson et al., 2013; Cichy et al., 2014; Contini et al., 2017; Grootswagers et al., 2019; Isik et al., 2013) and single unit recordings (e.g., Hung, Kreiman, Poggio, & DiCarlo, 2005). Behavioural data, including the reaction times collected from our participants, show that colour influences object identification speed (e.g., Bramão, Faísca, Petersson, & Reis, 2010). The neural data, however, did not show an effect of object colour on the classifier’s performance when distinguishing the neural activation patterns evoked by different object exemplars. For example, the brain activation pattern in response to a strawberry could be differentiated from the pattern evoked by a lemon, regardless of the congruency of their colours. This finding is consistent with previous results (Proverbio et al., 2004) but might seem puzzling because colour congruency has been shown to have a strong effect on object naming (e.g., Chiou et al., 2014; Nagai & Yokosawa, 2003; Tanaka & Presnell, 1999). One plausible possibility is that the source of behavioural congruency effects may be at the stage of response selection, which would not show up in these early neural signals. More evidence is needed, but there is no evidence in the current data to suggest colour congruency influences early stages of object processing.

Our data demonstrate that object representations are influenced by object-colour knowledge but not at the initial stages of visual processes. Consistent with a traditional hierarchical view, we show that visual object features are processed before the features are bound into a coherent object that can be compared with existing, conceptual object representations. However, our data also suggest that the temporal dynamics of colour processing are influenced by the typicality of object-colour pairings. Building on the extensive literature on visual perception, these results provide a timecourse for the integration of incoming visual information with stored knowledge, a process that is critical for interpreting the visual world around us.

## ACKNOWLEDGEMENTS

This research was supported by the Australian Research Council (ARC) Centre of Excellence in Cognition and its Disorders, International Macquarie University Research Training Program Scholarships to LT & TG, an ARC Future Fellowship (FT120100816) and an ARC Discovery project (DP160101300) to TC. ANR has funding from the ARC (DP12102835 and DP170101840). GLQ was supported by a joint initiative between the University of Louvain and the Marie Curie Actions of the European Commission [grant no: F211800012], with additional funding from the European Union’s Horizon 2020 research and innovation programme under the Marie Sklodowska-Curie grant agreement No 841909.

The authors acknowledge the University of Sydney HPC service for providing High Performance Computing resources.

